# Hybrid Systems Modeling for (Cancer) Systems Biology

**DOI:** 10.1101/035022

**Authors:** Roel Dobbe, Claire J. Tomlin

## Abstract

The advent of biological data of increasingly higher resolution in space and time has triggered the use of dynamic models to explain and predict the evolution of biological systems over space and time. Computer-aided system modeling and analysis in biology has led to many new discoveries and explanations that would otherwise be intractable to articulate without the available data and computing power. Nevertheless, the complexity in biology still challenges many labs in capturing studied phenomena in models that are tractable and simple enough to analyze. Moreover, the popular use of ordinary differential equation models have their limitations in that they solely capture continuous dynamics, while we observe many discrete dynamic phenomena in biology such as gene switching or mutations. Hybrid systems modeling provides a framework in which both continuous and discrete dynamics can be simulated and analyzed. Moreover, it provides techniques to develop approximations and abstractions of complex dynamics that are tractable to analyze.

## I. Motivation

In biology, many of the dynamic processes that we are interested in studying or affecting with treatments are inherently *complex*, i.e. nonlinear and varying across time and/or space. In the study of how biological systems behave we can make one important distinction between different dynamic phenomena, i.e. the difference between continuous and discrete dynamics. *Continuous state variables* model processes that evolve in some continuum, either time and/or space. This can be the evolution of protein expression over time or its diffusion throughout space. *Discrete state variables* model “sudden” changes or events in a system, such as a binary switch turning on or off, or, more involved, for a cancer cell switching through a sequence of distinct phenotypic states.

*Ordinary differential equations (ODEs) modeling* is generally excepted to be able to capture many physical and biological phenomena quantitatively as we observe them in nature and experiments. On the other hand, logical models like *Boolean networks (BNs)* seek completely qualitative rather than quantitative models of biological systems. BNs can succeed in capturing high-level discrete phenomena such as activation or deactivation with fewer parameters than their ODE counterpart and can be used to evaluate model structure. However, they cannot capture transient response, only steady state. Unfortunately, ODEs strict use of only continuous state variables is not able to model the discrete dynamics in a system, and vice-versa logical models cannot describe more complicated dynamical evolutions. This motivates the use of *hybrid system models*, that can capture both continuous dynamics and discrete events. Additionally, trying to capture dynamics that are nonlinear and varying in time or space with one ODE model can lead to expressions that are intractable to simulate and hard to analyze. Instead, one can often get away with modeling an *abstraction* with simple ODE models that approximate the dynamics locally in space or time, and use discrete state variables to model the connection between local models, yielding a hybrid system model that represents the overall system behavior. As such, the motivation for hybrid systems is two-fold: modeling continuous and discrete state dynamics in an integrated fashion, and building approximations that allow for tractable analysis, simulation and hypothesis development and testing.

## II. Modeling Framework

A dynamical system describes the evolution of state variables, typically real valued over time. Some dynamical systems can also be influenced by *exogenous inputs*, which represent either uncontrollable disturbances or controlled input signals. For example, we might be able to specifically target a certain protein on the cell membrane with an inhibitor, which is an input to the internal signaling pathway system. Other dynamical systems have *outputs*, which represent either quantities that can be measured (e.g. a biomarker), or quantities that need to be controlled or regulated (e.g. blood sugar level). Dynamical systems with both inputs and outputs are sometimes referred to as *control systems*. Based on the type of their state variables, dynamical systems may be classified into the following categories:

1. **Continuous State:** If the state takes values in Euclidean space ℝ*^n^* for some *n* ≥ 1. We will use *x* ∈ ℝ*^n^* to denote the state of a continuous dynamical system. This can for instance be the expression of a protein, or the proportion of cells with a specific phenotype.
2. **Discrete State:** If the state takes values in a finite or countable set {*q*_1_, *q*_2_,...}. We will use *q* to denote the state of a discrete system. Consider for example a cell which can switch between three distinct phenotypic states: basal, luminal or stem-like. In this case the discrete state denotes *q* ∈ {basal, luminal, stem-like}.
3. **Hybrid State Variables:** If a part of the state takes values in ℝ*^n^*, and another part takes values in a finite set. For example, consider the example of basal and luminal cells, each having their own “mode” that yields a unique signaling pathway dynamics - this is a hybrid system: part of the state (protein expressions) is continuous, while another part (namely the phenotypic state) is discrete.

To formalize the transition between discrete states of a hybrid systems model entail so-called edges, and a guard and reset function. Edges capture to which discrete states we can transition from each discrete state. The guard function models where in time *t* or space *x* a discrete state transition takes place or is triggered. The reset function explains what happens to the continuous state *x* as we go through a discrete state transition from *q_i_* to *q_j_*.

## III. Applications in Systems Biology

Since the application of hybrid systems modeling is relatively new to systems biology as a whole, and under-explored within cancer systems modeling, in this section, we provide examples of hybrid systems used for study of different biological systems. We distinguish between four important types of analysis and give examples for each.

- **Understanding the dynamic evolution of complex biological system:** It can be extremely tedious to do this for a system that has both continuous and discrete states. Moreover, most biomolecular systems of interest involve many interactions connected through positive and negative feedback loops for which an understanding of dynamics is hard to obtain. Simulating a hybrid system model can reveal insight into counterintuitive qualitative behaviors, that would otherwise be hard to interpret from experiments. In [1], the authors motivate the use of hybrid models for analyzing glucose metabolism in humans and spore formation in Bacillus subtilis. They argue that often encountered sigmoidal curves can be approximated by piecewise linear functions. In [2], our group studied the biologically observed equilibria of the multiple cell Delta-Notch protein signalling using reachability analysis, in which protein concentration dynamics inside each biological cell are modeled using linear differential equations; inputs activate or deactivate these continuous dynamics through discrete switches, which themselves are controlled by protein concentrations reaching given thresholds.
- **Studying multi-cellular ensembles:** The authors in [3] study multicellular behavior in prokaryotes that rely on cell-density-dependent gene expression. In Vibrio fischeri, a single cell is able to sense when a quorum of bacteria, a minimum population unit, is achieved. Under these conditions, certain behavior is efficiently performed by the quorum, such as bioluminescence. This is naturally modeled as a multi-modal hybrid system, resulting in simulations that are in accordance with experimental observations. In [4], our lab studied a multicellular pattern called Planar cell polarity (PCP) signaling in Drosophila melanogaster wings. PCP signalling generates subcellular asymmetry called autonomy, and through a poorly understood mechanism, mutant cell clones cause polarity disruptions of neighboring, wildtype cells, a phenomenon referred to as domineering non-autonomy. A cell-to-cell contact dependent signaling hypothesis, derived from experimental results is used to develop a hybrid system partial differential equation model. The sufficiency of this model, and the experimental validation of model predictions, reveal how specific protein-protein interactions produce autonomy or domineering non-autonomy. [5] provides a comprehensive survey of the use of hybrid models for tumor growth. They motivate the use of hybrid models to capture the multiscale nature of cancer and to handle multiple intra- and extracellular factors acting on different time and space scales. The cell centric nature of hybrid models naturally connects with cell biology and makes it possible to incorporate microenviromental components. Moreover, intracellular changes that result from mutations, altered intra- and intercellular signaling or protein trafficking can also be captured using hybrid models.
- **Parameter estimation and system identification:** In general, data-driven modeling of biological systems remains a difficult problem. The complexity and dimensionality of the modeled system often leads to ill-posed/underdetermined problems, and nonlinearities in system dynamics and measurement noise make this problem even more challenging. A key strategy to overcome these challenges is to use hybrid systems modeling to form higher-level abstractions, e.g. with compositions of simpler (often locally linear) models, that can grasp some low-dimensional structure of the system, while still representing biologically meaningful features that are worth analyzing. In [6], the authors study the problem of reconstructing dynamic fluxes and enzyme kinetics, using piecewise affine approximation models. The parameter estimation procedure is improved by separating it into two different phases. First, a dynamic flux profile in time is reconstructed using functions that are piecewise affine. Second, the time-dependent profiles are embedded in the concentration space and the enzyme kinetic functions for the single reactions are identified independently. The technique enables comparing different kinetic hypotheses very efficiently and thus promises to improve the biological knowledge of in vivo enzyme kinetics.
- **Modeling and identifying different network configurations with distinct dynamics:** Many cellular networks are capable of changing their configuration, through mutations or time-sequenced switching. If hypotheses for the different configurations and discrete events exist, hybrid models can be used to capture such phenomena. In [7], the authors study the problem of genetic network structure identification in a stochastic hybrid modeling framework. They considered a piecewise deterministic model of genetic networks where protein synthesis is triggered by discrete random binding events and follows simple deterministic kinetics. Using this approximation, they introduced an identification procedure that is based on matching the covariance function of the model to the data, which provides an estimate of the average effect of each transcription factor on every gene. In [8], a hybrid Boolean model (ODE+Boolean) is proposed for capturing biological signal pathways with postulated epigenomic feedback. The basic idea in this model is to combine continuous dynamical systems (an ODE model for already well-known parts of the network) with a discrete transition system (Boolean, for postulated but largely unknown components). The authors use an existing, well-known ODE model to “trigger” signal pathways represented by a Boolean model. This framework is easier to validate than a complete ODE model for large and complex signal pathways, for example to find unknown pathways to match the response to experimental data. The advantage of using a Boolean model for the unknown parts of the network is that relatively few parameters are needed. Thus, the framework avoids over-fitting, covers a broad range of pathways and easily represents various experimental conditions.

